# Estimating brain age from structural MRI and MEG data: Insights from dimensionality reduction techniques

**DOI:** 10.1101/859660

**Authors:** Alba Xifra-Porxas, Arna Ghosh, Georgios D. Mitsis, Marie-Hélène Boudrias

## Abstract

Brain age prediction studies aim at reliably estimating the difference between the chronological age of an individual and their predicted age based on neuroimaging data, which has been proposed as an informative measure of disease and cognitive decline. As most previous studies relied exclusively on magnetic resonance imaging (MRI) data, we hereby investigate whether combining structural MRI with functional magnetoencephalography (MEG) information improves age prediction using a large cohort of healthy subjects (*N*=613, age 18-88 yrs) from the Cam-CAN repository. To this end, we examined the performance of dimensionality reduction and multivariate associative techniques, namely Principal Component Analysis (PCA) and Canonical Correlation Analysis (CCA), to tackle the high dimensionality of neuroimaging data. Using MEG features (mean absolute error (MAE) of 9.60 yrs) yielded worse performance when compared to using MRI features (MAE of 5.33 yrs), but a stacking model combining both feature sets improved age prediction performance (MAE of 4.88 yrs). Furthermore, we found that PCA resulted in inferior performance, whereas CCA in conjunction with Gaussian process regression models yielded the best prediction performance. Notably, CCA allowed us to visualize the features that significantly contributed to brain age prediction. We found that MRI features from subcortical structures were more reliable age predictors than cortical features, and that spectral MEG measures were more reliable than connectivity metrics. Our results provide an insight into the underlying processes that are reflective of brain aging, yielding promise for the identification of reliable biomarkers of neurodegenerative diseases that emerge later during the lifespan.

## 1 Introduction

The human brain changes continuously across the adult lifespan. This process, termed brain aging, underlies the gradual decline in cognitive performance observed with aging. Although aging-induced changes are not necessarily pathological, the risk of developing neurodegenerative disorders rises with increasing age (Abbott, 2011), and diseases such as Alzheimer’s disease are thought to arise partly as a result of pathological processes associated with accelerated brain aging (Sluimer et al., 2009). Therefore, a better understanding of the neural correlates underlying brain aging, as well as better ways to identify biomarkers of healthy aging could contribute to improve the detection of early-stage neurodegeneration or predict age-related cognitive decline.

One promising approach for identifying individual differences in brain aging relies on the use of neuroimaging data to accurately predict “brain age” – the biological age of an individual’s brain (Cole et al., 2019b). In that context, machine learning (ML) techniques have proven to be a promising tool to ‘learn’ a correspondence between patterns in structural or functional brain features and the age of an individual (Dosenbach et al., 2010; Franke et al., 2010). ML techniques typically utilize functions in a high-dimensional space, whereby each dimension corresponds to a feature derived from neuroimaging data, to estimate the brain age. When predictive models are trained on neuroimaging datasets across the lifespan with a large number of subjects, they can generalize sufficiently well on unseen or ‘novel’ individuals. This provides the opportunity to deploy ML models at the population level and use the predicted age as a biomarker for atypical brain aging processes.

Most studies have explored the use of ML on data obtained from neuroimaging techniques to quantify atypical brain development in diseased populations. A common practice entails training a ML-based prediction model on healthy subjects and subsequently using it to estimate brain age in patients. The difference between an individual’s predicted brain age and their chronological age is then computed (the “brain age delta”), providing a potential measure that indicates increased risk of pathological changes that may lead to neurodegenerative diseases. For instance, this approach has been applied to study brain disorders and diseases including Alzheimer’s disease (Franke and Gaser, 2012; Gaser et al., 2013), traumatic brain injury (Cole et al., 2015), schizophrenia (Koutsouleris et al., 2014; Schnack et al., 2016; Shahab et al., 2019), epilepsy (Pardoe et al., 2017), dementia (Wang et al., 2019), Down’s syndrome (Cole et al., 2017a), Prader-Willi syndrome (Azor et al., 2019), and several others (Kaufmann et al., 2019), as well as other pathologies such as chronic pain (Cruz-Almeida et al., 2019), HIV (Cole et al., 2017c) and diabetes (Franke et al., 2013). Additionally, brain age prediction has also been extended beyond understanding neurological disorders such as in the context of testing the positive influence of meditation (Luders et al., 2016), as well as education and physical exercise (Steffener et al., 2016b) on brain age. Recent work has also shown a relationship between the brain age delta and specific cognitive functions, namely visual attention, cognitive flexibility, and semantic verbal fluency (Boyle et al., 2020).

The studies mentioned above have mainly focused on estimating brain age based on structural magnetic resonance imaging (MRI), with most studies using T1-weighted images (e.g. Cole, Leech and Sharp, 2015; Cole, Poudel, *et al.*, 2017). This is partly due to the availability of large lifespan MR-based open datasets, which has allowed researchers to train and validate their predictive models on a large number of subjects. However, it is well known that in addition to structural alterations, changes in brain function also occur during aging (Cabeza et al., 2018; Grady, 2012; Peters, 2006). One example of brain function changes associated with age is functional connectivity, which measures the statistical interdependence between time series from distinct brain regions (Sala-Llonch et al., 2015). Functional connectivity measures derived from functional MRI (fMRI) data have been successfully used to predict age (Dosenbach et al., 2010; Li et al., 2018; Nielsen et al., 2018; Vergun et al., 2013). However, the improvement of brain age prediction and ultimately the detection of early disease stages using fMRI is limited due to the sluggishness of the hemodynamic response function, which severely limits the time resolution at which the underlying neural events can be resolved and consequently the ability to perform directional connectivity analysis (Smith et al., 2011). Abnormal synchronization processes at faster timescales have been detected in several neuropsychiatric disorders (Schnitzler and Gross, 2005), and in particular in movement disorders such as Parkinson’s disease (Heinrichs-Graham et al., 2014; Kondylis et al., 2016; Moisello et al., 2015; Nelson et al., 2017) and essential tremor (Kondylis et al., 2016; Raethjen and Deuschl, 2012; Schnitzler et al., 2009). Therefore, neuroimaging techniques with higher temporal resolution, such as electroencephalography (EEG) and magnetoencephalography (MEG), can offer complementary features associated with both normal and pathological aging. In particular, MEG provides higher spatiotemporal resolution compared to EEG because magnetic fields propagate with little attenuation and distortion (Baillet, 2017). Recently, several studies have investigated age-related brain function changes using EEG (Dimitriadis and Salis, 2017; Sun et al., 2019; Zoubi et al., 2018), which has enabled researchers to build a brain age prediction model based on the temporal and spectral features of electrophysiological brain activity, as well as the connectivity between brain regions. A detailed overview of different neuroimaging modalities and ML methods that have been used to estimate brain age is presented in (Cole et al., 2019a).

The aforementioned studies investigated the age-related structural and functional brain changes separately. Recently, several researchers have examined the combination of features from different modalities and demonstrated that this could lead to better brain age prediction performance. For instance, Liem and colleagues combined anatomical features extracted from MR images and fMRI connectivity measures (Liem et al., 2017). Another recent study combined MRI anatomical features, fMRI connectivity measures and MEG features (Engemann et al., 2020). However, changes in brain tissue composition with age alter the T1 signal intensity of brain structures (Cho et al., 1997; Salat et al., 2009), possibly as a result of changes in iron concentration (Harder et al., 2008; Ogg and Steen, 1998). Studies using MR-derived anatomical features such as cortical thickness and subcortical volume are thus not taking into account age-related changes in tissue signal properties that may improve brain age prediction performance. Therefore, combining whole brain T1 signal intensities with functional (EEG/MEG or fMRI) features yields potential to this end. Further research is needed to determine the extent to which this approach can improve the latter.

Moreover, a major roadblock to clinical applications of ML models is their explainability (Bzdok and Ioannidis, 2019), in other words the ability to attribute their predictions to specific input variables. From a clinical perspective, it would be useful to identify the neuroimaging features that are important to the ML model to estimate brain age. As argued by Kriegeskorte *et al*., decoding models can reveal whether information pertaining to a specific outcome or behavioural measure is present in a particular brain region or feature (Kriegeskorte and Douglas, 2019). In the same study, the authors also highlighted the difficulties and confounds associated with interpreting weights in a linear decoding model and consequently suggested the use of multivariate techniques to identify the most informative predictors. The current brain age prediction models (e.g. Azor et al., 2019; Cole et al., 2017b) have used a similarity metric to retrieve lower dimensional embeddings from neuroimaging features. However, this technique does not warrant explainability, as it relies on the neuroimaging features similarity between individuals to make predictions. Therefore, it is of interest to explore dimensionality reduction techniques that allow identification of the informative neuroimaging features for age prediction.

To address the aforementioned challenges related to the combined use of structural and functional neuroimaging data to predict age and the explainability of the associated ML models, the main aims of the present study were to: (i) investigate whether adding functional information from MEG recordings to the whole-brain MRI voxel intensity features would improve brain age prediction performance, (ii) examine the performance of dimensionality reduction techniques in conjunction with ML models, and (iii) improve the explainability of ML-based brain age prediction by applying multivariate associative statistical methods for identifying key features that exhibit the most prominent age-related changes. To do so, we used structural MRI and functional MEG data collected from a large cohort of healthy subjects. We applied Principal Component Analysis (PCA) and Canonical Correlation Analysis (CCA) as dimensionality reduction and multivariate associative techniques, respectively, to assess their predictive performance compared to the widely used similarity metric technique. Finally, we identified and visualized the most informative features in the context of age prediction.

## 2 Materials and Methods

### 2.1 Dataset

We analyzed data from the open-access Cambridge Center for Aging Neuroscience (Cam-CAN) repository (see Shafto et al. 2014; Taylor et al. 2017 for details of the dataset and acquisition protocols), available at https://camcan-archive.mrc-cbu.cam.ac.uk//dataaccess/. Specifically, we used structural (T1-weighted MRI) and functional (resting-state MEG) neuroimaging data from 652 healthy subjects (male/female = 322/330, mean age = 54.3 ± 18.6, age range 18-88 years). The MR images were acquired from a 3T Siemens TIM Trio scanner with a 32-channel head coil. The images were acquired using a MPRAGE sequence with TR = 2250 ms, TE = 2.99 ms, Flip angle = 9°, Field of View = 256 × 240 × 192 mm^3^ and voxel size = 1 mm isotropic. The resting-state MEG data were recorded using a 306-channel Elekta Neuromag Vectorview (102 magnetometers and 204 planar gradiometers) at a sampling rate of 1kHz. For the resting-state scan, subjects were asked to lie still and remain awake with their eyes closed for around 9 min. Following exclusions (e.g. subjects that did not have both MRI and MEG data, unsatisfactory pre-processing results such as failure to remove cardiac and ocular artifacts and/or failure to extract the cortical surface for source reconstruction), we report findings from a final dataset including 613 subjects. A descriptive list of subjects included in our dataset is detailed in the Supplementary Materials, and Supp. Fig. 1 depicts gender and age distributions for the included participants.

### 2.2 Neuroimaging data processing

A summary of the feature extraction process for MR images and MEG recordings is illustrated in Fig. 1.

**Figure 1.**
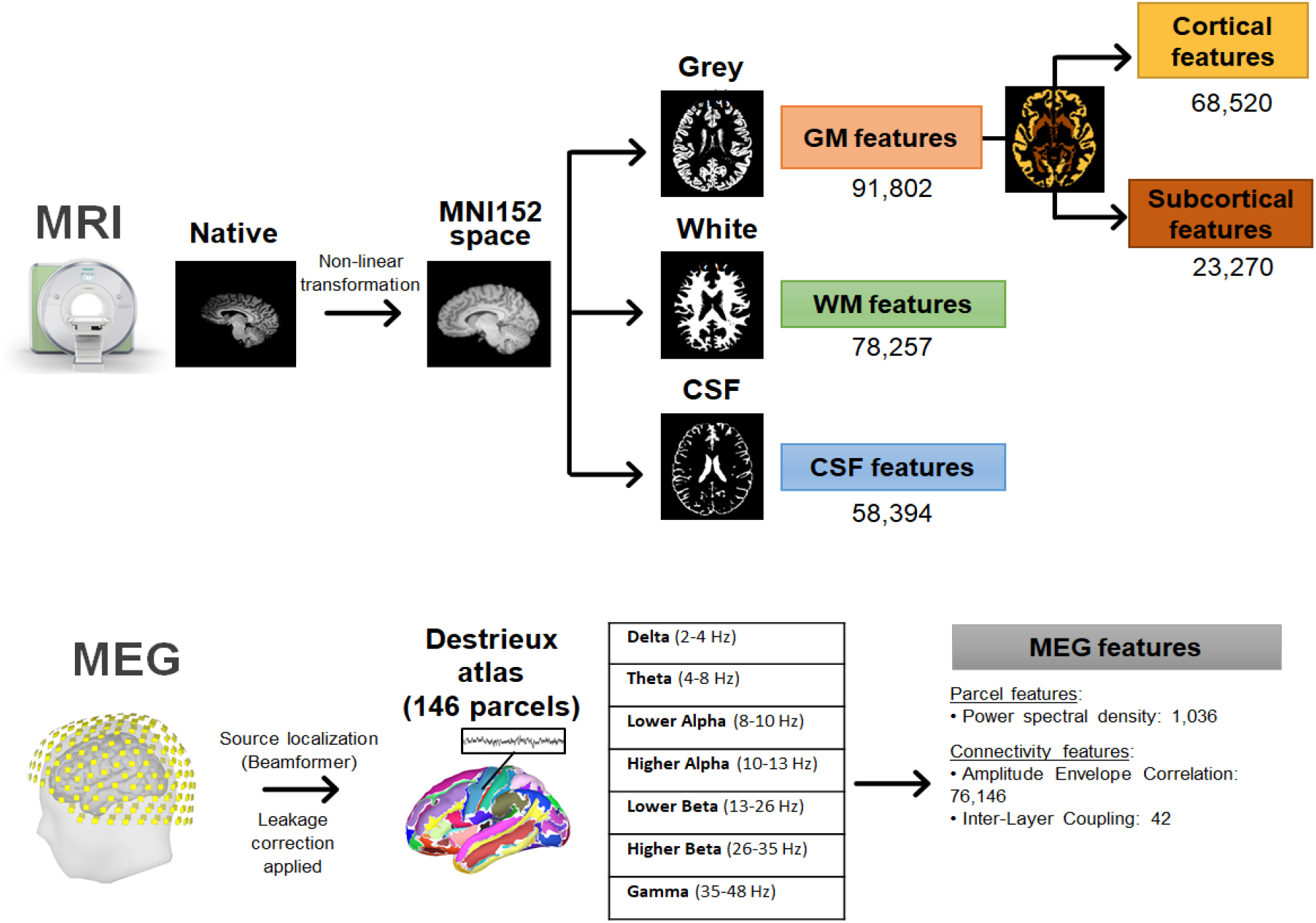
Feature extraction pipeline for MRI and MEG data.

#### 2.2.1 MRI structural analysis

The processing of T1-weighted MR images followed the pipeline presented in (Cole et al., 2017b) and was implemented using tools from the FMRIB Software Library (FSL, http://www.fmrib.ox.ac.uk/fsl) (Jenkinson et al., 2012). Briefly, the Brain Extraction Tool (BET) (Smith, 2002) was used to isolate the brain tissue, and the FMRIB’s Linear/Nonlinear Image Registration Tools (FLIRT/FNIRT) (Andersson et al., 2007; Jenkinson and Smith, 2001) were used to perform a non-linear registration to the MNI152 template brain (2mm resolution). Next, the registered images were segmented into Grey Matter (GM), White Matter (WM), and Cerebrospinal Fluid (CSF) using the MNI152 template mask for each tissue type. The GM maps were further segmented into cortical and subcortical regions to delineate the effects of aging on these regions. The resultant images were vectorized and subsequently z-scored to obtain a feature vector for each subject. This process resulted in a feature matrix where each row consisted of normalized intensity values for a single subject (see Fig. 1 for the exact number of features from each brain structure).

#### 2.2.2 MEG analysis

The MEG data were processed using the open-source software MNE-Python (https://martinos.org/mne) (Gramfort et al., 2014). Raw MEG data were high-pass filtered at 1 Hz, notch filtered at 50 Hz and 100 Hz to remove power line artifacts, and resampled at 200 Hz. Cardiac and eye movement artifacts were identified using Independent Component Analysis (ICA) and automatically classified comparing the ICA components with the simultaneously recorded electrocardiography (ECG) and electrooculography (EOG) signals (Jas et al., 2018). Artifact-free MEG data were converted from sensor to source space on the subject’s cortical surface using the linearly constrained minimum variance (LCMV) beamformer (Van Veen et al., 1997). The cortical surface was reconstructed from the T1-weighted MR images as obtained from the FreeSurfer recon-all algorithm (Dale et al., 1999; Fischl et al., 2004, 2002, 2001, 1999a, 1999b; Fischl and Dale, 2000). The sources were constrained within the cortical regions of the brain and assumed to be perpendicular to the cortical envelope. The noise covariance matrix was estimated using the empty room recordings, and the data covariance matrix was estimated directly from the MEG data. After source reconstruction, we parcellated the cortex into 148 brain regions using the Destrieux atlas (Destrieux et al., 2010). Each parcel time series were corrected for signal leakage effects using a symmetric, multivariate correction method intended for all-to-all functional connectivity analysis (Colclough et al., 2015). For each parcel, the power spectral density (PSD) for the entire resting state scan was calculated and averaged within 7 frequency bands, namely Delta (2–4 Hz), Theta (4–8 Hz), lower Alpha (8–10 Hz), higher Alpha (10-13 Hz), lower Beta (13–26 Hz), higher Beta (26–35 Hz) and Gamma (35–48 Hz). Relative power was calculated by dividing the power within each band by the total power across all bands (Niso et al., 2019). In addition to the PSD values, amplitude envelope correlation (AEC) within each frequency band was used to estimate the functional connectivity between different cortical parcels (Brookes et al., 2012; Hipp et al., 2012), as this method provides a robust measure for stationary connectivity estimation (Colclough et al., 2016). Inter-layer Coupling (ILC) was also calculated from the functional connectivity matrices to estimate the similarity of the connectivity profile across frequency bands (Tewarie et al., 2016). Therefore, each row of the resulting MEG feature matrix consisted of PSD, AEC and ILC values for a single subject (see Fig. 1 for the exact number of features from each type).

### 2.3 Brain age prediction analysis

We examined the performance of three dimensionality reduction techniques: the similarity metric, PCA and CCA. The MRI and MEG features were embedded into a lower dimensional space using these techniques. The transformed features were subsequently used as input features in the prediction models. The higher dimensional feature set prior to transformation was also directly used as input to evaluate the potential improvement achieved by using dimensionality reduction techniques. All the prediction models were implemented using the scikit-learn toolbox (Pedregosa et al., 2011) in Python.

#### 2.3.1 Dimensionality reduction techniques

##### 2.3.1.1 Similarity metric

Following the approach presented by Cole and colleagues (Cole et al., 2017b), we represented the data as a *N* × *N* similarity matrix (*N* being the number of subjects in the training set). The similarity between any two subjects was calculated using the dot product between their corresponding feature vectors. Therefore, each testing subject was represented as an *N*-element vector containing similarity values corresponding to each of the *N* training subjects.

However, the use of a similarity metric entails the following issues: (1) the training set requires an adequate number of subjects to sample the spectrum of healthy aging completely, and (2) the predictions are based on how similar a test subject is to each of the training subjects. To address these issues, we used dimensionality reduction techniques, namely PCA and CCA, to identify the neuroimaging features that mostly contribute to brain age prediction. Specifically, PCA and CCA project the data onto a lower dimensional space and allow ML models to represent age as a function of neuroimaging features, as opposed to similarity scores. This approach allowed us to visualize the age-related neuroimaging features after the model was trained.

##### 2.3.1.2 Principal Component Analysis (PCA)

PCA is a linear dimensionality reduction technique using singular value decomposition (SVD) of the data to project it onto a lower dimensional space (Jolliffe, 2002). It is widely used to decompose multivariate datasets into a set of successive orthogonal components that explain the maximum amount of the data variance (e.g. Amico and Goñi, 2017; Larivière *et al.*, 2019). The obtained principal components correspond to the maximal modes of variation and hence correspond to the most prominently changing features in the dataset. Often, the number of principal components is selected visually as the point where the total variance explained by increasing the number of components starts plateauing (“knee rule”). We applied PCA to project the feature matrix onto a lower dimensional space and subsequently estimate brain age using a prediction model. In each case, we used the knee of the curve relating the variance explained vs the number of principal components to decide the number of components to be retained. In all the models using only MRI data, 5 components were retained, whereas in all models using MEG data or a combination of MRI and MEG data, 10 components were retained. The number of retained principal components explained about 60-66% of the variance in the data.

##### 2.3.1.3 Canonical Correlation Analysis (CCA)

CCA is an alternative dimensionality reduction technique that identifies latent variables to model the covariance between some input and output variables (Thompson, 2005). CCA has been successfully applied in the context of brain-behavior relationships (Smith et al., 2015), neurodegenerative diseases (Avants et al., 2014) and psychopathology (Xia et al., 2018). CCA, similarly to PCA, uses the SVD factorization method to reduce the dimensionality of the data. However, in CCA the covariance matrix is used instead of the input variance matrix. Therefore, the obtained canonical components are maximally correlated to the output variable.

In the present case, the CCA inputs were the neuroimaging feature matrices and the output the chronological age vector. Therefore, CCA retrieved a linear combination of the neuroimaging features that were maximally correlated to the age of the subjects. We used CCA to project the feature vector along this direction and subsequently used the projection values to predict age.

CCA yields a loading vector for every CCA component, which quantifies the contribution of each feature to that CCA component (Wang et al., 2018). We used these loading values to assess the contribution of each feature to brain age prediction and thereby understand which regions of the brain mostly exhibited age-related changes. To estimate the reliability of these loading values, we used the bootstrapped ratio, whereby we repeated the CCA analysis for 1000 bootstrapped samples of the dataset chosen at random with replacement (Efron and Tibshirani, 1986; McIntosh and Lobaugh, 2004). The bootstrapped ratio (BSR) of the loading values indicates which areas reliably contribute to the brain age prediction, thus increasing the overall reliability of the prediction models. The procedure for generating the BSR of the loading values is illustrated in Fig. 2.

**Figure 2.**
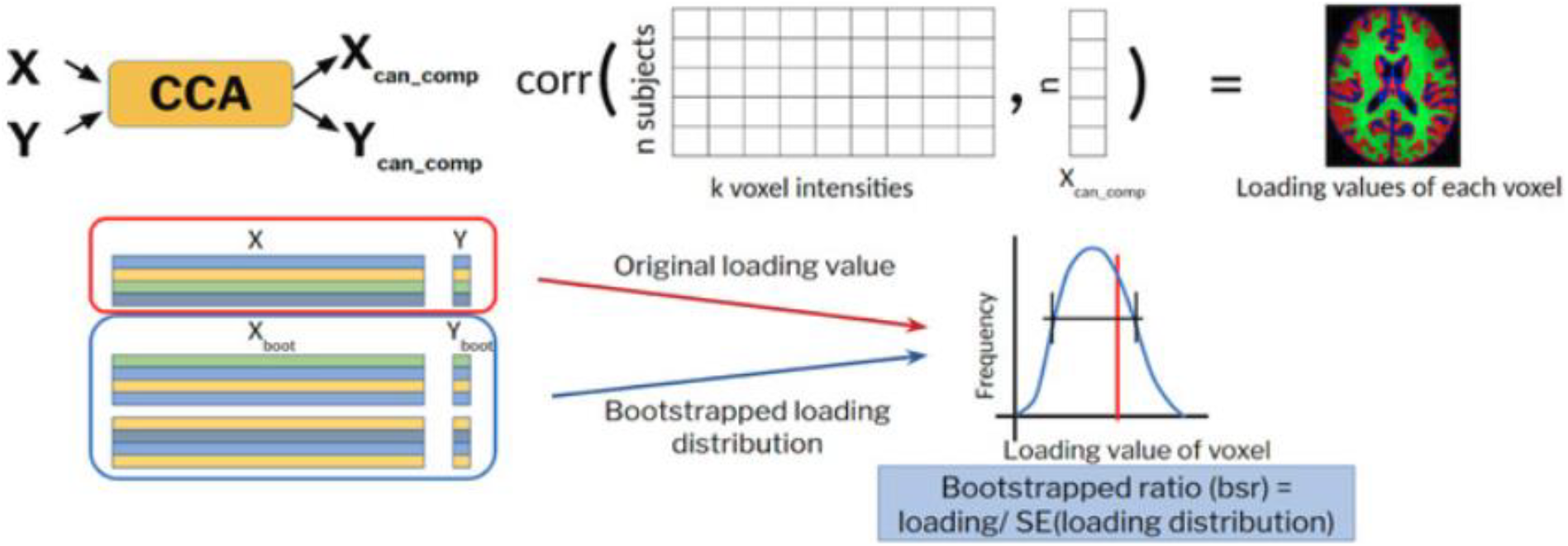
Calculation of loadings and bootstrapped ratio (BSR) of loading values from the employed CCA model.

We also examined deep CCA (Andrew et al., 2013) to learn a non-linear combination of features that maximally covary with age. However, deep CCA was not numerically stable and hence it was not explored further.

#### 2.3.2 Prediction model

Predictive models using MRI or MEG features independently were implemented using Gaussian Process Regression (GPR) models with an additive dot-product and white kernel. The GPR models were defined using the neuroimaging features as inputs (i.e. independent variables) and chronological age as the output (i.e. dependent variable). GPR has been widely used for predicting chronological age from T1-weighted images (Aycheh et al., 2018; Cole et al., 2018, 2017a, 2017b, 2017c, 2015). GPR is a non-parametric approach, which finds a distribution over possible functions that are consistent with the data (Rasmussen and Williams, 2006). The main assumption underlying GPR is that any finite subset of the available data must follow a multivariate Gaussian distribution. The prior belief about the relationship between variables is decided by the sufficient statistics of these multivariate Gaussian distributions, namely the mean vector and standard deviation matrix. The standard deviation matrix, therefore, indicates the confidence of model predictions. Multivariate Gaussian distributions also have the ability to reflect local patterns of covariance between individual data points. Therefore, a combination of multiple such distributions in a Gaussian process can model non-linear relationships and is more flexible than conventional parametric models, which rely on fitting global models. The implemented GPR models contained two hyperparameters that required tuning, the inhomogeneity of the dot-product and the noise level of the white kernel, which were selected among five candidate values ranging between 0.01 and 100.

Models combining both MRI and MEG features were implemented through a stacking framework using random forest regression, following recent age prediction studies proposing model-stacking strategies to combine features from different neuroimaging modalities (Engemann et al., 2020; Liem et al., 2017). To aggregate the information from MRI and MEG data, a feature vector was constructed, which comprised the cross-validation predictions from the single-modality GPR models and the corresponding uncertainty in prediction (characterized by the standard deviation of GPR prediction). This feature vector contained four features per subject and was used to train the random forest model. Note that the same training-testing splits as in the single-modality models were used to train the random forests to ensure they were tested on left-out predictions. The random forests contained two tunable hyperparameters: the number of trees, which was set to 10, 50, or 100, and the tree-depth, which was set to 5, 10, 20 or None (None indicating splitting till leaf nodes contained only one sample).

The performance of each model was evaluated using a nested 10-fold cross-validation strategy and scored based on the mean absolute error (MAE) between estimated and chronological age. The hyperparameter selection for the GPR and random forest models was done by grid search within a 5-fold inner loop. Specifically, we performed the dimensionality reduction followed by fitting the regression model on the training set of each fold of the inner loop. We repeated these steps for multiple sets of hyperparameter values and selected the hyperparameter set that yielded the best performance across the validation sets of the inner loop. Using this optimal hyperparameter set, we repeated the dimensionality reduction and fitting the regression model on the training set of the outer fold. This pipeline was thereafter applied on the test set to evaluate its performance on new data points. The nested cross-validation scheme was repeated 10 times to attain a less biased estimate of model performance, hence 100 MAE values were obtained for each model. To get an estimate of the chance level of age prediction, we used predictions from a random model with no training. Irrespective of the modality of data used, the chance level of MAE was ~16.74 years and *R*^2^ was around zero. These values served as a baseline to assess the performance of the examined prediction models.

The codes implementing all the preprocessing and examined prediction models are available on GitHub at https://github.com/axifra/BrainAge_MRI-MEG.

## 3 Results

We compared the performance of all models using a 10-fold cross-validation approach and repeated the cross-validation framework 10 times. A summary of the performance of the different brain age prediction methods is presented in Fig. 3, in terms of the MAE difference with respect to an MRI-only model using GPR. In Table 1, we report the absolute MAE values. All models, irrespective of the data modality, performed better than chance level (MAE ~16.74 years) thus indicating that all the considered neuroimaging features exhibited some age-related effects.

**Table 1.** Comparison of age prediction by GPR models combined with different dimensionality reduction techniques based on MRI and MEG features, as well as stacking of both modalities. Mean absolute error (MAE) values were calculated over the testing set (mean **±**standard deviation).

| Input Features | Model | MAE (years) | R<sup>2</sup> |
| --- | --- | --- | --- |
| MRI | GPR | 5.88 ± 0.56 | 0.83 ± 0.05 |
| | Similarity+GPR | 5.61 ± 0.72 | 0.85 ± 0.05 |
| | PCA+GPR | 8.07 ± 0.68 | 0.69 ± 0.07 |
| | CCA+GPR | 5.33 ± 0.51 | 0.86 ± 0.03 |
| MEG | GPR | 9.60 ± 0.98 | 0.55 ± 0.09 |
| | Similarity+GPR | 9.80 ± 0.94 | 0.54 ± 0.10 |
| | PCA+GPR | 12.64 ± 1.12 | 0.28 ± 0.13 |
| | CCA+GPR | 9.68 ± 0.89 | 0.55 ± 0.09 |
| MRI+MEG (Stacking) | GPR | 4.97 ± 0.53 | 0.88 ± 0.03 |
| | Similarity+GPR | 5.17 ± 0.60 | 0.87 ± 0.04 |
| | PCA+GPR | 7.26 ± 0.73 | 0.73 ± 0.07 |
|  | CCA+GPR | **4.88 ± 0.52** | **0.88 ± 0.03** |

**Figure 3.**
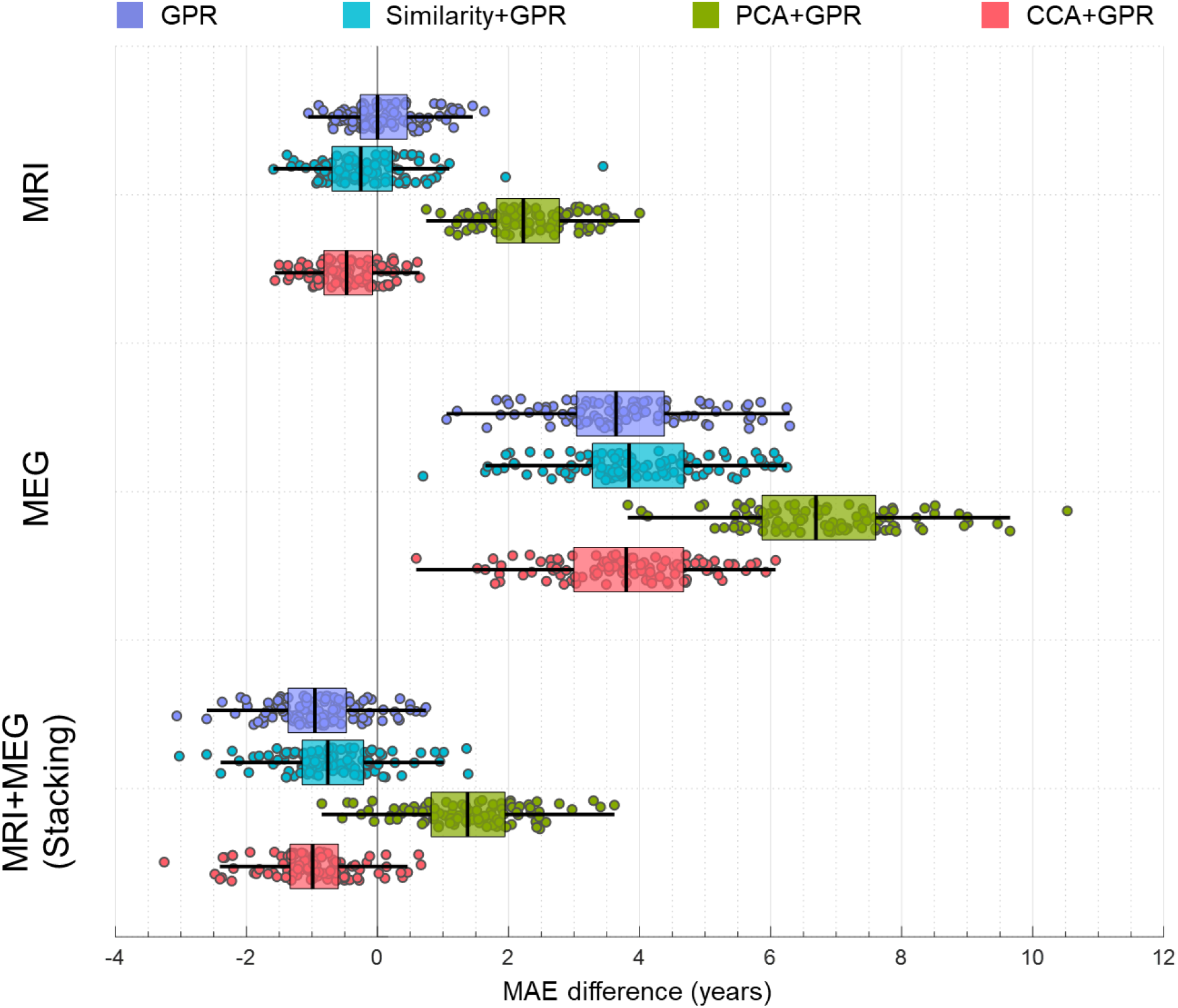
Boxplots depicting the age prediction performance for models using different dimensionality reduction techniques based on MRI and MEG features, as well as combining both data modalities using a stacking model. Each dot represents the mean absolute error (MAE) difference from the MRI-based GPR model at a given fold (10 k-folds × 10 repetitions). The best performance was obtained with the stacking model, either using GPR alone or GPR combined with CCA, showing an improvement of around 1 year with respect to the MRI-based GPR model. PCA degraded performance in all cases.

### 3.1 Dimensionality reduction techniques

#### 3.1.1 MRI features

The voxel-wise T1-weighted intensity levels (from all tissues) were used as input to the different dimensionality reduction techniques. CCA+GPR yielded the best performance with respect to age prediction, with a corresponding MAE of 5.33 years (Table 1), which corresponded to an improvement of around 0.5 years compared to feeding the high dimensional feature vector directly into a GPR model, and an improvement of around 0.3 years compared to the similarity metric method (Fig. 3). PCA resulted in a significantly inferior performance, yielding a MAE of 8.07 years (Table 1). The failure of the similarity metric to yield the best performance is likely due to the sample size of the dataset, which was much smaller than the dataset used in (Cole et al., 2017b). This could possibly have led to incomplete sampling of the aging subspace and hence yielded worse performance.

To delineate the contribution of cortical and subcortical MRI features, we compared the performance for each of these features separately. Subcortical MRI features clearly outperformed cortical features, irrespective of the model used (Fig. 4, Supp. Table 1). The GPR model and CCA+GPR method yielded similar performance, with subcortical MRI features yielding a MAE of ~5.76 years and cortical MRI features yielding a MAE of ~7.11 years. The performance of the similarity metric technique was slightly worse and PCA, as before, resulted in a significantly poorer performance compared to all other techniques. These results indicate that the subcortical regions were more reliable indicators of brain age compared to cortical regions. This finding was further supported by the CCA loadings of MRI features, whereby subcortical regions exhibited higher BSR of loading values compared to cortical regions (Fig. 6b).

**Figure 4.**
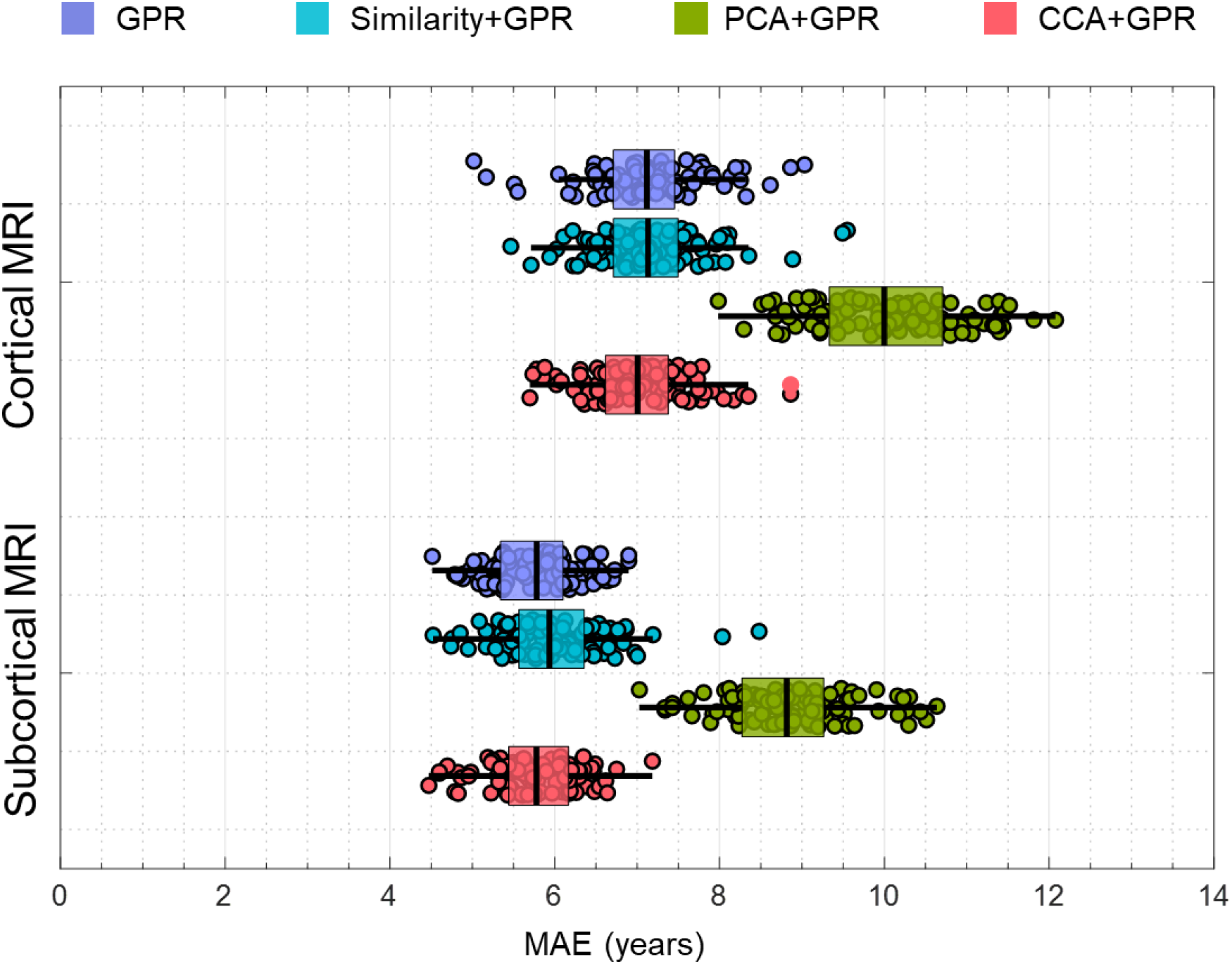
Boxplots depicting the age prediction performance for models using either cortical or subcortical MRI features, using different dimensionality reduction techniques. Each dot represents the mean absolute error (MAE) at a given fold (10 k-folds × 10 repetitions). Better performance was obtained with subcortical MRI features compared to cortical MRI features.

#### 3.1.2 MEG features

The MEG features extracted from the source-space MEG data were the relative PSD within seven frequency bands for each brain region examined, as well as the AEC and ILC measures to quantify functional connectivity (Tewarie et al., 2016). Using the high dimensional feature vector directly as the input to a GPR model yielded similar performance as the models transforming the features to a low dimensional embedding using the similarity metric or CCA, with a MAE of ~ 9.70 years (Table 1). On the other hand, PCA yielded inferior performance, similarly to the MRI features (Table 1). Overall, MEG features yielded a considerably inferior performance compared to MRI features (Fig. 3).

An important consideration when comparing MEG age prediction performance to MRI is that the MEG features only contained information from the cortex, whereas the MRI intensities were from both cortical and subcortical regions. Therefore, we compared the performance of models using MEG features to those including only cortical MRI features. MEG features still yielded worse performance for all the examined methods, compared to using cortical MRI features (worse by 2.49 years for GPR, 2.66 years for similarity, 2.59 years for PCA, and 2.67 years for CCA respectively). This suggests that MEG features were worse predictors of brain age than MRI features.

We subsequently explored which MEG features were better at predicting brain age. We found that ILC values did not significantly contribute to age prediction, with the corresponding MAE values being very close to those of the random model (Supp. Table 2). PSD performed better than AEC when using the high dimensional feature vector directly into a GPR model, but AEC performed better than PSD when using the CCA+GPR model (Supp. Table 2).

### 3.2 Combining structural and functional features from MRI and MEG data

We next examined the potential benefit of combining MEG and MRI features compared to using MRI features alone. Note that we included MRI features from all structural tissues (GM, WM and CSF) and both spectral and connectivity MEG features. To build the multimodal prediction, we combined the age predictions from each modality using a stacking model. All models exhibited improved performance when both modalities were used for age prediction (Table 1). Without dimensionality reduction, we found that using the stacking model yielded an age prediction improvement of around one year compared to using MRI-only features (Fig. 3). Furthermore, CCA yielded the best performance with a MAE of 4.88 years (Table 1), improving the performance of CCA+GPR model using MRI-only features by around 0.5 years. These results suggest that functional MEG features contained complementary information to anatomical MRI features, thereby providing non-redundant information that improved the estimation of brain age.

Recently, several studies have reported an age-related bias in estimates of brain age, commonly observed as an overestimation of age in younger subjects and an underestimation of age in older subjects (Aycheh et al., 2018; Cole and Franke, 2017; Liang et al., 2019; Smith et al., 2019). Our cohort had less subjects within the lower and higher age ranges (Supp. Fig. 1), which could have led to an age-related bias in the estimates. To investigate this, in Fig. 5 we show the predicted vs. chronological age for the best model using both MRI and MEG features (stacking CCA+GPR). The fitted model yields a good match with the ideal model, and no bias is observed for the youngest and oldest subjects.

**Figure 5.**
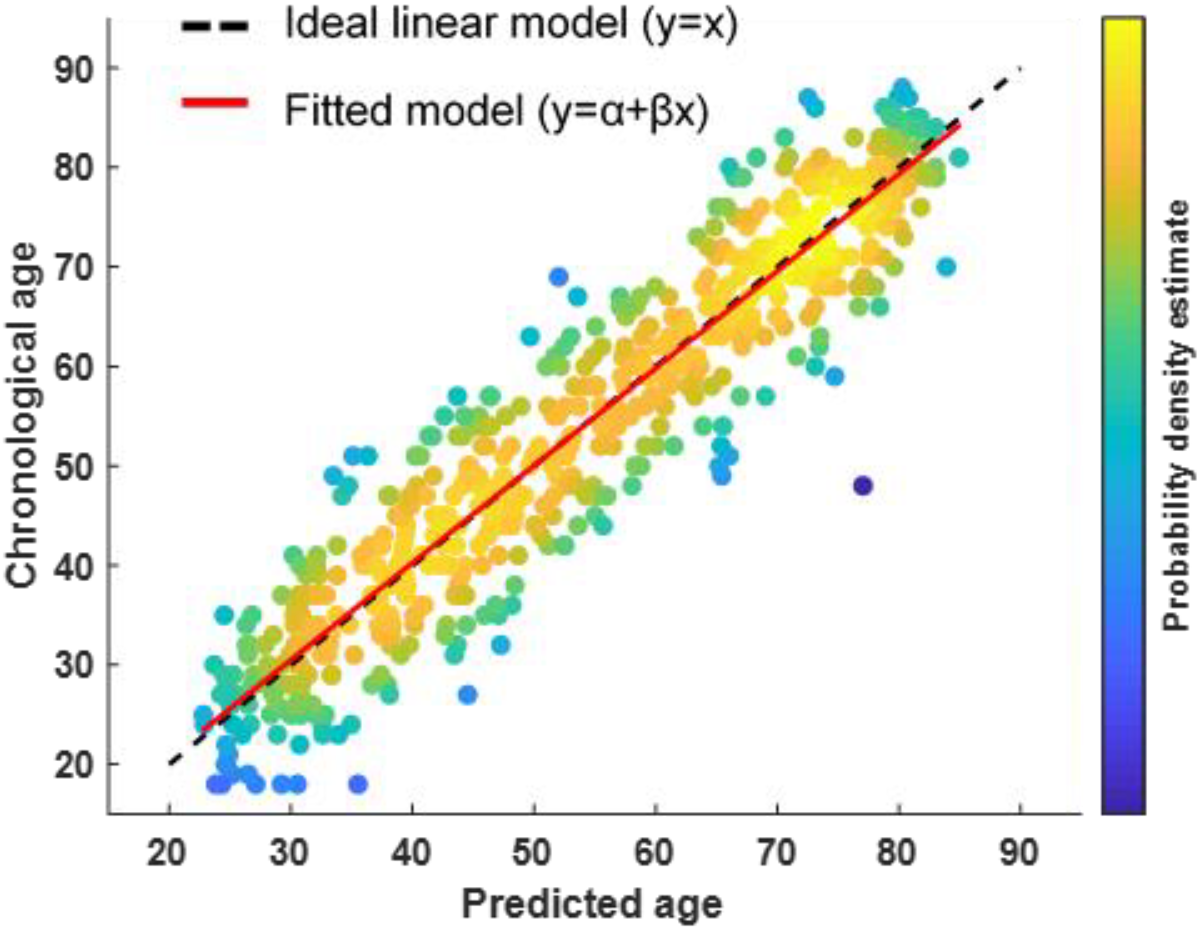
Plot of predicted vs. chronological age for the stacking CCA+GPR model using both MRI and MEG features. The black dotted line corresponds to the ideal linear model (***y*** = ***x***), whereas the red line corresponds to the fitted model (***y*** = ***α*** + ***βx***).

### 3.3 CCA loadings

One of the goals of the present work was to identify the brain regions exhibiting more pronounced age-related changes. The CCA loadings provided a way to assess the contribution of each neuroimaging feature to age prediction, thus indicating the features that yielded the most reliable age-related changes. The histogram of the BSR of voxel intensity loading values, as well as the top 15% BSR of loading values for GM, WM, and CSF are shown in Fig. 6a & 6b, respectively. The histogram of BSR values indicates that GM and WM voxels exhibited more reliable age-related changes as compared to CSF (the histogram peak for GM and WM was located around −300 and −400 respectively, whereas the histogram peak for CSF was located around −100). Almost all of the loading values were negative, indicating a decreased voxel intensity with increasing age. Further, the top 15% of BSR values corresponded to subcortical regions, thus supporting our results that these regions yield better age prediction (Supp. Table 1). Some of these areas are shown in Fig. 6b, while 3D nifti volumes of the CCA loadings are available in NeuroVault (https://identifiers.org/neurovault.collection:6091). The highlighted GM areas were localized in subcortical structures such as the putamen, thalamus, and the caudate nucleus, as well as regions in the cerebellum. Most of the highlighted WM voxels were confined to the corpus callosum, thus indicating that the latter was associated to the most consistent age-related changes among WM voxels. Another structure among WM voxels that exhibited age-related changes was the thalamic radiation.

**Figure 6.**
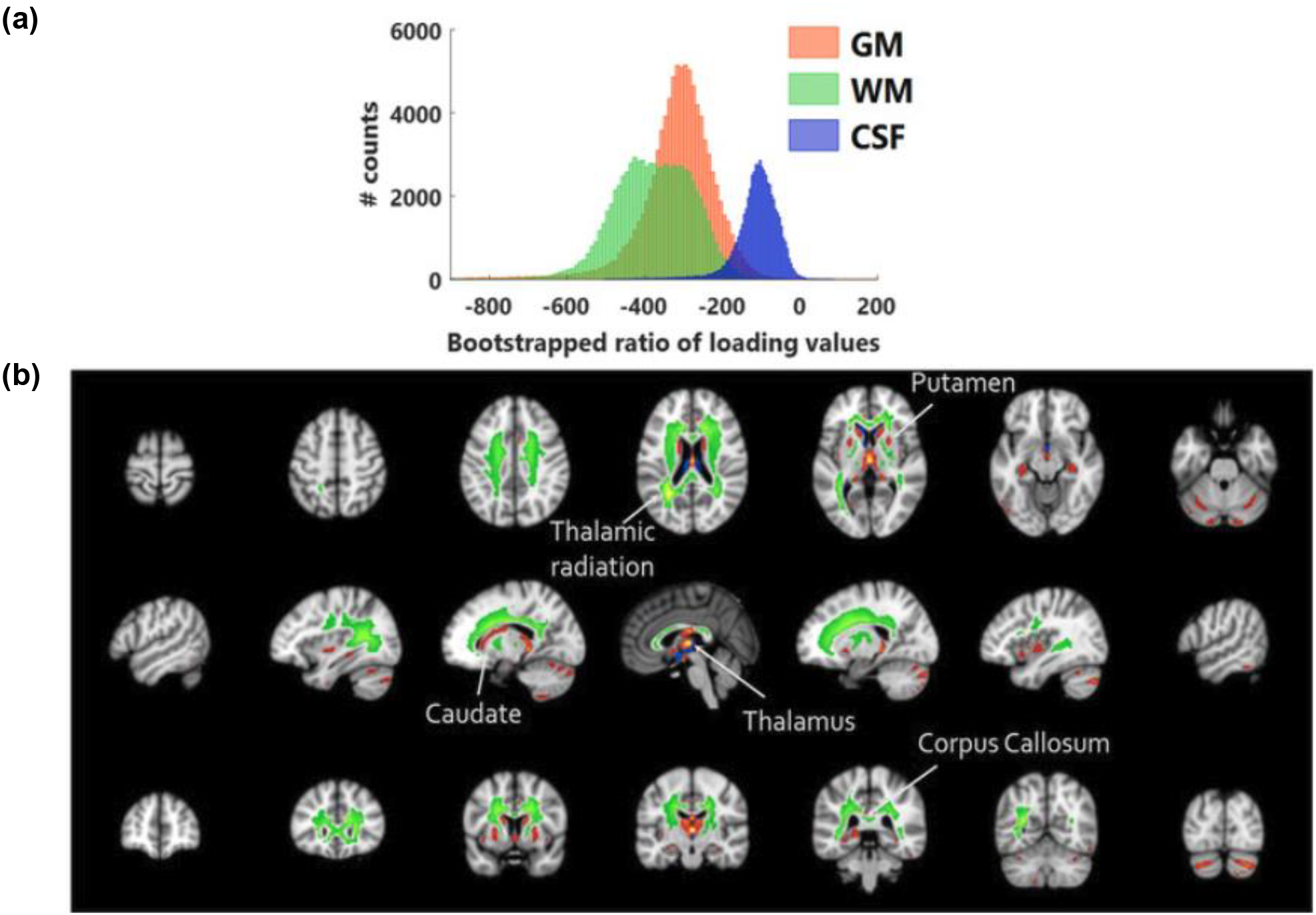
BSR of CCA loading values for T1-weighted intensity levels. (a) Distribution of BSR values for GM (orange), WM (green) and CSF (blue) voxels. (b) Brain regions with the top 15% BSR values, highlighting that the most reliable voxels for brain age prediction were located within subcortical regions.

Furthermore, we visualized the CCA loadings for the MEG features. The PSD loadings are shown in Fig. 7 and the AEC loadings are shown in Supp. Fig. 2. We found that PSD values were more reliable (BSR values ~450) than AEC values (BSR values ~300). Regarding the PSD loadings (Fig. 7), we observed different regions showing age-related effects within various frequency bands. Contrary to MRI loadings, whereby most of the loading values were negative, PSD loadings were found to be both positive and negative. The low-frequency bands exhibited decreasing PSD values with age, with delta and theta band PSD exhibiting maximal age-related effects in the frontal areas and alpha band PSD exhibiting maximal age-related effects in the visual and motor areas. Higher frequency bands (beta and gamma) exhibited increasing PSD values with age in frontal and motor areas. Regarding the AEC loadings (Supp. Fig. 2), the all-to-all connectivity matrices (one per frequency band) were sorted by functional networks according to the Yeo 7-network brain cortical parcellation (Yeo et al., 2011). Most functional connections exhibited increased connectivity with age within all frequency bands, with the exception being the visual network, which exhibited decreased connectivity with age for the high alpha and high beta frequency bands. The ILC loadings are not shown since ILC values did not significantly contribute to age prediction (Supp. Table 2).

**Figure 7.**
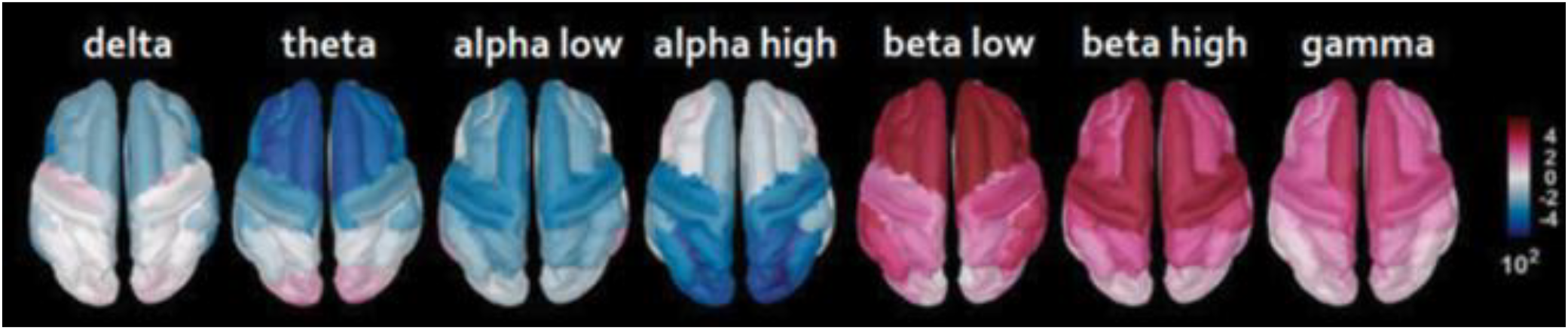
BSR of CCA loading values depicting cortical regions with PSD values that are correlated (pink) or anti-correlated (blue) with age for each frequency band.

## 4 Discussion

In this study, we aimed to leverage multimodal neuroimaging data to predict brain age in a large cohort of healthy subjects (*N*=613) between 18-88 years and assess the performance of several dimensionality reduction techniques. We found that applying a CCA+GPR model to each imaging modality and combining their predictions through a stacking model yielded the best performance (Fig. 3), with a MAE of 4.88 years. Conversely, we found that PCA yielded inferior performance for brain age prediction, regardless of the imaging modality (Fig. 3). Furthermore, we identified and visualized the regions that exhibited age-related changes and found that subcortical T1-weighted intensity levels were more informative for age prediction compared to cortical regions (Figs. 4, 6). We also identified age-related changes in the spectral features of various cortical regions using MEG data (Fig. 7). In addition, we demonstrated that using multivariate associative techniques such as CCA yields better explainability of the predictive models, which may contribute to the identification of clinically relevant biomarkers of pathological aging.

### 4.1 Dimensionality reduction techniques

We used T1-weighted MR images and resting-state MEG data to develop a brain-age prediction framework that uses both structural and functional information of the brain. We restricted our analysis to cortical sources of the MEG data and thereby had functional information from cortical regions only. Since the goal was to predict age, the desired MAE for the perfect model would be 0 years. However, owing to subject variability and the ill-conditioning of the problem, specifically the definition of a “healthy” subject and the assumption that chronological age should perfectly correspond to brain age, we did not expect to achieve a MAE of 0 years.

In the present study we used GPR as the regression model of choice for brain age prediction. Furthermore, we explored the contribution of dimensionality reduction techniques to age prediction. A commonly used dimensionality reduction technique in neuroimaging studies is PCA. However, PCA yielded inferior prediction performance for both imaging modalities (Fig. 3). This result may be explained by the fact that large variability exists between neuroimaging features across subjects. As argued by (Stringer et al., 2019), PCA decomposition yields a lower dimensional space where the data manifold is smoother compared to the original data manifold in the high-dimensional neuroimaging feature space. Given that this smoother manifold is comprised of the top principal components, it is indicative of the low frequency information in the data. Our results indicate that this lower dimensional principal component-induced manifold is not predictive of brain age, thereby implying that aging causes higher frequency changes in the original data manifold. The high frequency nature of the age-related information suggests that neighbouring points (each point representing a subject) on the data manifold may have different brain age values. As stated earlier, similarity between neuroimaging features of two subjects could be guided by other phenotypical factors, thus resulting in the two subjects from different age groups being placed in each other’s neighbourhood on the data manifold. Additionally, PCA yields components that are maximally varying in the dataset, which could be aligned to directions of subject variability in the dataset instead of age-related changes. Therefore, our results suggest that using PCA to perform dimensionality reduction does not lead to good performance in the context of brain age prediction. In contrast, CCA improved performance by yielding the component that maximally covaries with age, therefore identifying features that are informative for age prediction.

### 4.2 Combining structural and functional features from MRI and MEG data

Combining anatomical information from MR images and functional information from MEG recordings resulted in an improvement in brain age prediction for all models (Fig. 3). Particularly, the performance of the GPR model with the high dimensional MRI features improved by around one year when the MEG features were also considered, and the CCA+GPR model performance improved by around 0.5 years. The superior performance of the age prediction model when combining structural and functional features suggests that both modalities carry complementary information that are to some degree independent. Our results are in agreement with a recent study that used MRI and MEG data from the CamCAN dataset to estimate brain age where it was reported that combining both modalities showed an improvement in age prediction of around 0.8 years compared to MRI-only prediction (Engemann et al., 2020). Comparing the age prediction results in absolute values, Engemann et al. reported a MAE of 5.2 years, whereas our models achieved better performance (GPR model: 4.97 years, CCA+GPR: model 4.88 years). A key difference between the two studies is that the MRI features considered by Engemann *et al*. were cortical thickness, cortical surface area and subcortical volume, whereas in this study we used whole-brain MRI voxel intensity features. Therefore, our results suggest that brain age prediction models may benefit from exploiting the rich information contained in MR images, instead of extracting specific anatomical features from them.

### 4.3 CCA loadings

Apart from yielded the best prediction accuracy, CCA was used to identify the brain regions that contribute more reliably to age prediction. CCA returns loading values for each input feature, therefore improving model explainability. Using the BSR of loading values for MRI features, we found that most of the voxel T1-weighted intensity levels were negatively correlated with age (Fig. 6a). A decrease in voxel intensities with age has been reported by (Salat et al., 2009), who suggested that this association was an indicator of brain atrophy. Thus, our findings are in agreement with previous studies that have reported cortical thinning with age (Fjell et al., 2009; Hogstrom et al., 2013; Salat et al., 2004; Storsve et al., 2014).

Furthermore, our results indicate that subcortical regions are more reliable predictors of age compared to cortical regions. The brain structures that most reliably exhibited age-related changes included the putamen, thalamus, and caudate nucleus, which are important structures involved in relaying a variety of information across the brain, in sensorimotor coordination, and in higher cognitive functions (Grahn et al., 2008; Sefcsik et al., 2009; Sherman and Guillery, 2002). A number of stereological and MRI studies have reported atrophy in subcortical regions associated with aging, specifically in the putamen (Bugiani et al., 1978), amygdala (Coffey *et al.*, 1992; Fjell *et al.*, 2013), hippocampus (Fjell *et al.*, 2013; Nobis *et al.*, 2019), caudate nucleus (Krishnan et al., 1990), substantia nigra (McGeer et al., 1977), thalamus (Sullivan *et al.*, 2004; Fjell *et al.*, 2013), and cerebellum (Andersen et al., 2003; Good et al., 2001; Torvik et al., 1986). Recent studies using large subject cohorts have also reported an age-related decrease in the hippocampal and temporal lobe volumes (Nobis *et al.*, 2019). Hence, our findings are in agreement with the changes in size of specific brain areas associated with aging as reported in previous relevant studies. Furthermore, we explored whether there was an association between the decrease in volume for several subcortical regions and their respective BSR of CCA loadings, illustrated in Fig. 8. We found that a higher CCA loading was associated with a larger decrease in volume across age (R=0.44, *p*=0.08). This suggests that the CCA loadings, to some extent, reflect the shrinkage of subcortical structures. However, it is likely that they are also associated with increased iron deposition in subcortical areas with age (Harder et al., 2008; Ogg and Steen, 1998). Moreover, we cannot rule out the possibility that the absence of strong negative correlations between age and MRI voxel intensities in the cortex could be attributed to improper alignment of sulci and gyri to the standard MNI152 brain template.

**Figure 8.**
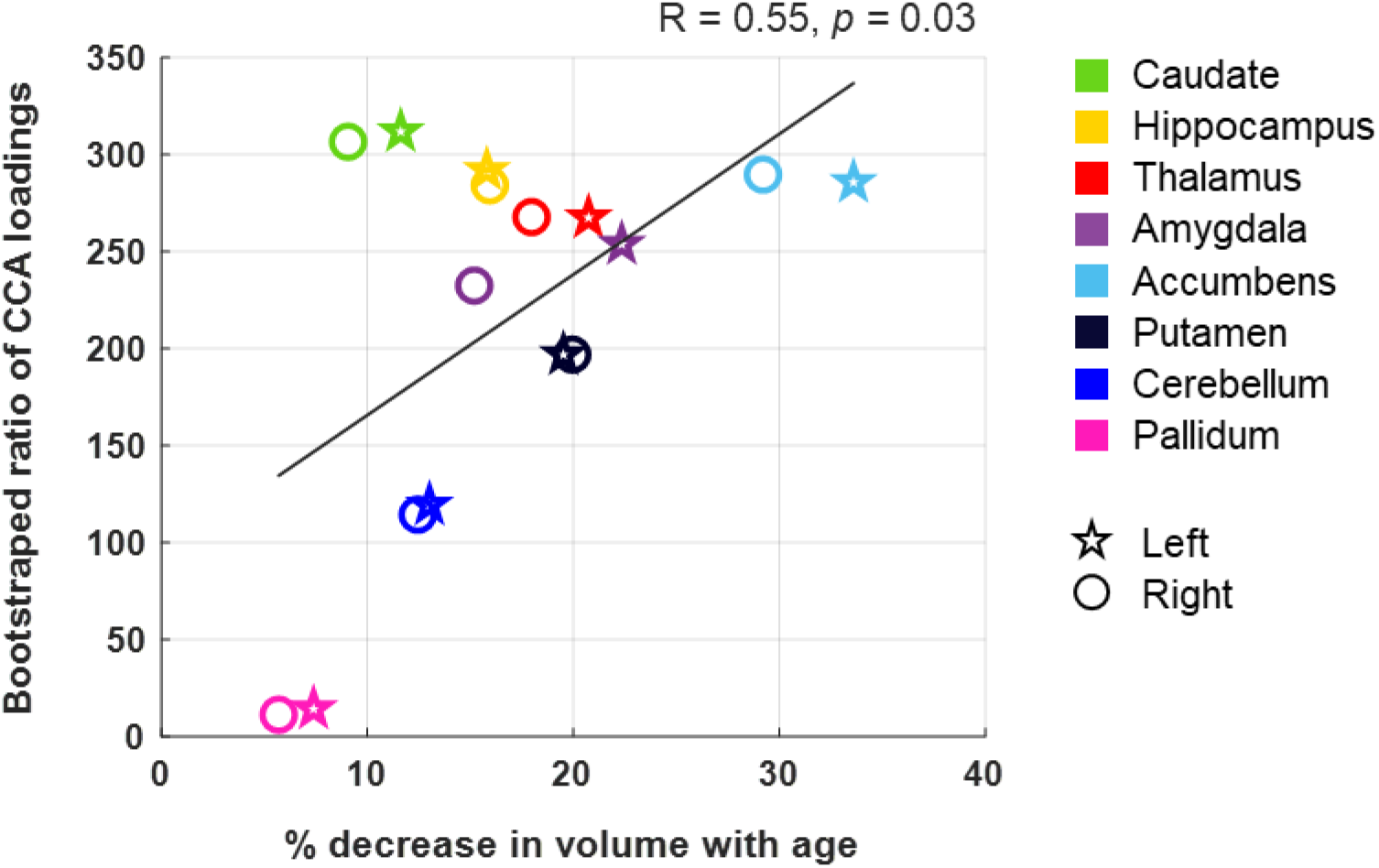
Association between the percentage decrease in volume with age and the bootstrapped ratio (BSR) of CCA loadings for each subcortical region. A significant correlation was observed, whereby a higher CCA loading was associated with a larger decrease in volume across age. The subcortical volume of each structure was extracted using FreeSurfer for each subject. Note that the BSR of CCA loadings are depicted as absolute values.

We found that the WM regions affected by age were mostly confined to the corpus callosum and the thalamic radiation. These results are in strong agreement with previous studies that have reported age-related alterations in WM structures (Salat et al., 2005), such as atrophy in corpus callosum fiber tracts (Ota et al., 2006; Pfefferbaum et al., 2000) and thalamic radiation (Cox *et al.*, 2016). Although CCA loadings for CSF voxels did not exhibit high BSR values compared to their GM and WM counterparts, including CSF improved model performance. CSF information possibly indicates changes in brain volume and ventricle size that resulted in improved brain age prediction.

Among the examined MEG features, PSD and AEC values yielded the best performance; however, the PSD values were found to be more reliable than AEC values (compare BSR values in Fig. 7 and Supp. Fig. 2). These results align with previous EEG and MEG studies (Dimitriadis and Salis, 2017; Engemann et al., 2020; Sun et al., 2019; Zoubi et al., 2018), which reported improved brain age prediction using power spectral features. We found the BSR of loading values for PSD values to be both positively and negatively correlated with age, depending on the frequency band (Fig. 7). Our results revealed that delta and theta power decreases with age, most prominently in frontal regions. These results are in agreement with the fact that slower waves (0.5–7 Hz) have been reported to decrease in power in older adults as compared to their younger counterparts (Caplan et al., 2015; Cummins and Finnigan, 2007; Leirer et al., 2011; Vlahou et al., 2014). Increased frontal theta activity has been linked to better performance in memory tasks (Jensen and Tesche, 2002; Onton et al., 2005), which may explain the decreasing power in lower frequencies for increasing age. Regarding the alpha band, the strongest effect of age was observed in the occipital cortex, whereby increased power within the higher alpha subband (10-13 Hz) was negatively correlated with age. These results align with several studies that have reported an association between a decrease in alpha power and increasing age (Gómez et al., 2013; Hübner et al., 2018). However other studies have not reported significant changes in alpha power with age (Heinrichs-Graham and Wilson, 2016; Xifra-Porxas et al., 2019). Likely, the later studies did not have sufficient statistical power to detect this age-related decrease in alpha power, since the cohort size was below 35 subjects, whereas the studies that reported an association between alpha power and age (including ours) had a sample size larger than 85. Nevertheless, it is worth pointing out that the observed reduced alpha power in older adults could be a result of dividing the alpha power in conventional lower and upper alpha bands, considering that a recent study reported that younger and older adults had equivalent alpha power at the individual alpha peak frequency (Scally et al., 2018).

In line with many previous studies, we observed an association between beta power and age (Heinrichs-Graham and Wilson, 2016; Hübner et al., 2018; Rossiter et al., 2014; Xifra-Porxas et al., 2019). Specifically, we found that the age-related increase in lower beta power (13-26 Hz) was restricted to frontal regions, whereas higher beta power (26-35 Hz) was restricted to the motor cortex. This beta power increase has been linked to higher levels of intracortical GABAergic inhibition as tested by pharmacological manipulations (Hall et al., 2011; Muthukumaraswamy et al., 2013). This suggests that the age-related changes in beta power may be associated with greater GABAergic inhibitory activity within motor cortices of older subjects.

Finally, we found that AEC measures exhibited an age-related increase in connectivity within all frequency bands across all brain networks, apart from the visual network which showed a decrease in connectivity within the high alpha and high beta frequency bands. These results align well with a recent study where Larivière et al. reported lower beta-band connectivity in the visual network and higher beta-band connectivity in all other brain networks with age (Larivière et al., 2019). Higher functional connectivity in older adults has been associated with a lower cognitive reserve (López et al., 2014), and individuals with mild cognitive impairment exhibit an enhancement of the strength of functional connections (Bajo et al., 2010; Buldú et al., 2011). Overall, the results from these studies suggest that the age-related increase in MEG functional connectivity, as seen in our study, may play a role in modulating cognitive resources to compensate for the lack of efficiency of the memory networks (Bajo et al., 2010), and therefore represent a marker of the decline in cognitive functions observed during aging.

### 4.4 Limitations

A limitation of brain age prediction is the use of chronological age as a surrogate for brain age. Although we used a cohort of healthy subjects, brain age is known to depend on various other factors, such as education (Steffener et al., 2016a). In this work, we ignored all lifestyle factors and aimed to predict the biological age from neuroimaging features. Furthermore, we used a single model to predict the brain age for both males and females. These factors contribute to the biological age labels being an imperfect surrogate of the “true” brain age of each subject.

Moreover, the MEG features extracted in this study were restricted to cortical regions. As MRI features from subcortical structures were found to be the best age predictors, we speculate that including functional features from deep brain structures could have resulted in greater improvement in the prediction models. This suggests that the use of newly developed methodologies to more reliably detect brain activity in deeper structures using MEG (Pizzo et al., 2019) could contribute to improved age prediction in future studies.

## 5 Conclusions

Leveraging structural and functional brain information from MRI and MEG data, we showed that combining features from both modalities using a stacking model yielded better performance compared to using a single neuroimaging modality. We showed that dimensionality reduction techniques can be used to improve brain age prediction and identify key neuroimaging features that reflect age-related effects. Specifically, we found that using CCA in conjunction with GPR yielded the best age prediction performance, whereas using PCA deteriorated prediction performance. We also showed that the most reliable MRI predictors of age-related effects were features derived from subcortical structures such as the putamen, thalamus, and caudate nucleus, and WM regions such as the corpus callosum. Finally, we found that spectral MEG features were more reliable than connectivity metrics.

## Supporting information

List of subject IDs

## Acknowledgments

We wish to thank Drs. Stefanie Blain-Moraes, Karim Jerbi and Danilo Bzdok for valuable discussions regarding this work. Data collection and sharing for this project was provided by the Cambridge Centre for Ageing and Neuroscience (Cam-CAN). Cam-CAN funding was provided by the UK Biotechnology and Biological Sciences Research Council (grant number BB/H008217/1), together with support from the Medical Research Council (MRC) Cognition & Brain Sciences Unit (CBU) and the European Union Horizon 2020 LifeBrain project. This work was supported by funds from the Natural Sciences and Engineering Research Council of Canada (NSERC) Discovery Grants RGPIN-2017-05270 [MHB] and RGPIN-2019-06638 [GDM], the Fonds de la Recherche du Quebec - Nature et Technologies (FRQNT) Team Grant 254680-2018 [GDM & MHB], and scholarships from McGill University and Quebec Bio-Imaging Network [AXP & AG], as well as and MITACS [AG]. The research was undertaken thanks in part to funding from the Canada First Research Excellence Fund, awarded to McGill University as part of the Healthy Brains for Healthy Lives initiative.

## Supplementary material

**Supp. Table 1.** Comparison of age prediction by GPR models combined with different dimensionality reduction techniques based on MRI features from grey matter (GM), white matter (WM), as well as cortical vs subcortical GM. Mean absolute error (MAE) values were calculated over the testing set (mean **±**standard deviation).

| Model | Input Features | MAE (years) |
| --- | --- | --- |
| GPR | GM | $6.12 \pm 0.61$ |
| | WM | $6.39 \pm 0.64$ |
| | Cortical GM | $7.11 \pm 0.68$ |
|  | **Subcortical GM** | **$5.76 \pm 0.54$** |
| Similarity + GPR | GM | $6.20 \pm 0.63$ |
| | WM | $6.56 \pm 0.66$ |
| | Cortical GM | $7.14 \pm 0.68$ |
| | Subcortical GM | $5.98 \pm 0.63$ |
| PCA + GPR | GM | $9.32 \pm 0.81$ |
| | WM | $10.98 \pm 1.07$ |
| | Cortical GM | $10.05 \pm 0.89$ |
| | Subcortical GM | $8.84 \pm 0.76$ |
| CCA + GPR | GM | $6.02 \pm 0.52$ |
| | WM | $6.38 \pm 0.54$ |
| | Cortical GM | $7.01 \pm 0.60$ |
|  | **Subcortical GM** | **$5.78 \pm 0.53$** |

**Supp. Table 2.**
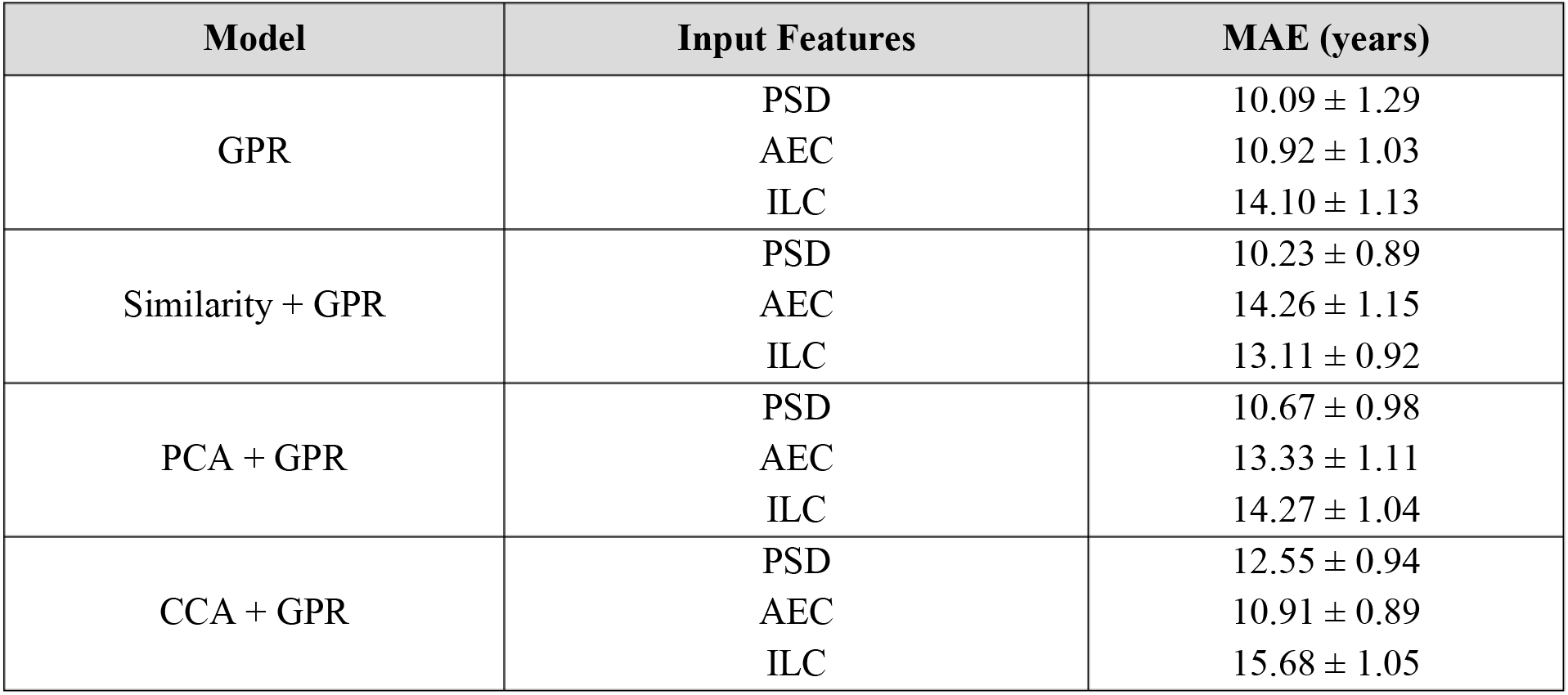
Comparison of age prediction by GPR models combined with different dimensionality reduction techniques based on MEG features. Mean absolute error (MAE) values were calculated over the testing set (mean **±**standard deviation).

| Model | Input Features | MAE (years) |
| --- | --- | --- |
| GPR | PSD | $10.09 \pm 1.29$ |
| | AEC | $10.92 \pm 1.03$ |
| | ILC | $14.10 \pm 1.13$ |
| Similarity + GPR | PSD | $10.23 \pm 0.89$ |
| | AEC | $14.26 \pm 1.15$ |
| | ILC | $13.11 \pm 0.92$ |
| PCA + GPR | PSD | $10.67 \pm 0.98$ |
| | AEC | $13.33 \pm 1.11$ |
| | ILC | $14.27 \pm 1.04$ |
| CCA + GPR | PSD | $12.55 \pm 0.94$ |
| | AEC | $10.91 \pm 0.89$ |
| | ILC | $15.68 \pm 1.05$ |

**Supp. Fig. 1.**
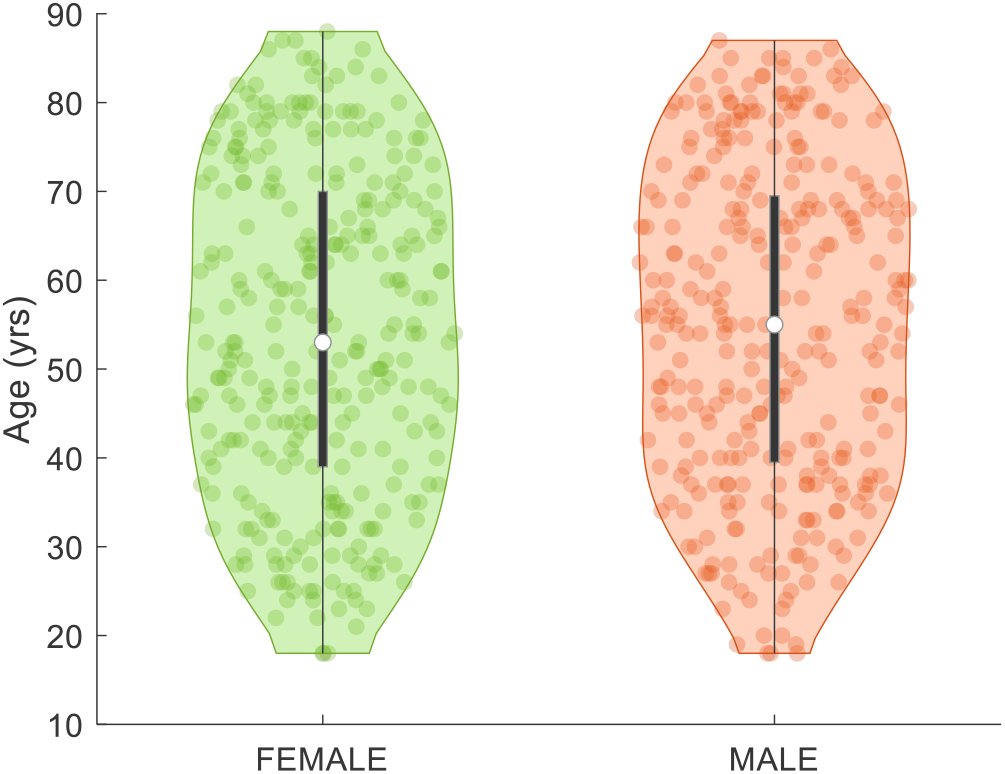
Demographic data for the included participants from the Cam-CAN dataset.

**Supp. Fig. 2.**
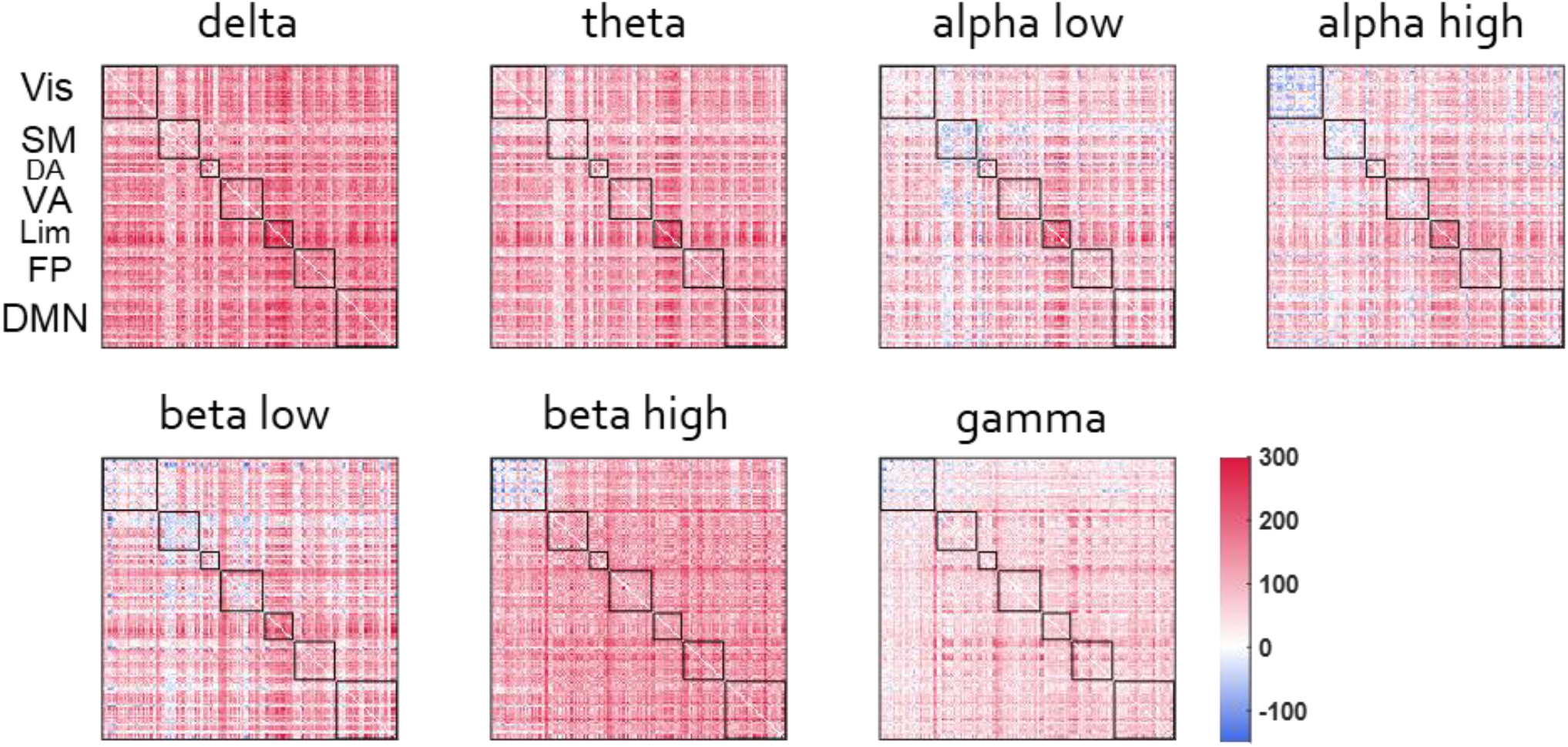
BSR of CCA loadings for AEC values depicting functional connections for which strength increased (red) or decreased (blue) with age for each frequency band. The AEC matrices were sorted by functional networks according to the Yeo 7-network brain cortical parcellation (Vis=Visual, SM=Somatomotor, DA=Dorsal attention, VA=Ventral attention, Lim=Limbic, FP=Frontoparietal, DMN=Default mode network). Most functional connections tended to increase their strength with age, except from the connectivity within the visual network for the alpha high and beta high frequency bands.

